# Ionic conductances driving tonic firing in Purkinje neurons of larval zebrafish

**DOI:** 10.1101/2023.11.29.569172

**Authors:** Meha P. Jadhav, Shivangi Verma, Vatsala Thirumalai

**Author notes:** These authors contributed equally to this work (Meha P. Jadhav, Shivangi Verma). Department of Biology, University of Konstanz, Konstanz 78464, Germany (Meha P. Jadhav).

## Abstract

Purkinje neurons are critical for the functioning of the cerebellum, which is among the oldest and most conserved regions of the vertebrate brain. In mammals and in larval zebrafish, Purkinje neurons can generate tonic firing even when isolated from the network. Here we investigated the ionic basis of tonic firing in Purkinje neurons of larval zebrafish using voltage clamp for isolation of membrane currents along with pharmacology. We discovered that these neurons express L-type and P/Q-type high voltage-gated calcium currents, T-type low voltage-gated calcium currents and SK and BK-type calcium dependent potassium currents. Among these, L-type calcium currents and SK-type calcium-dependent potassium currents were indispensable for tonic firing, while blocking T-type, P/Q-type and BK currents had little effect in comparison. We observed that action potentials were broadened when either L-type or SK channels were blocked. Based on these results, we propose that calcium entry via L-type calcium channels activates SK potassium channels leading to faster action potential repolarization, in turn aiding the removal of inactivation of sodium channels. This allows larval zebrafish Purkinje neurons to continue to fire tonically for sustained periods. In mammals also, tonic firing in Purkinje neurons is driven by calcium channels coupling to calcium-dependent potassium channels, yet the specific types of channels involved are different. We therefore suggest that coupling of calcium channels and calcium-dependent potassium channels could be a conserved mechanism for sustaining long bouts of high frequency firing.

**Key points:**

- Tonic firing is an intrinsic property of Purkinje neurons in mammals and fish.
- These neurons express multiple types of voltage-gated conductances including L-type, T-type, and P/Q-type calcium currents and SK- and BK-type calcium-dependent potassium currents.
- Blocking L-type calcium channels and SK-type calcium dependent potassium channels resulted in spike broadening and reduced tonic firing.
- L-type calcium currents were activated during the repolarization of the spike.
- Based on this we conclude that calcium entry via L-type channels activates SK-channels causing faster repolarization of the spike and therefore sustained tonic firing.

**Graphical abstract:** 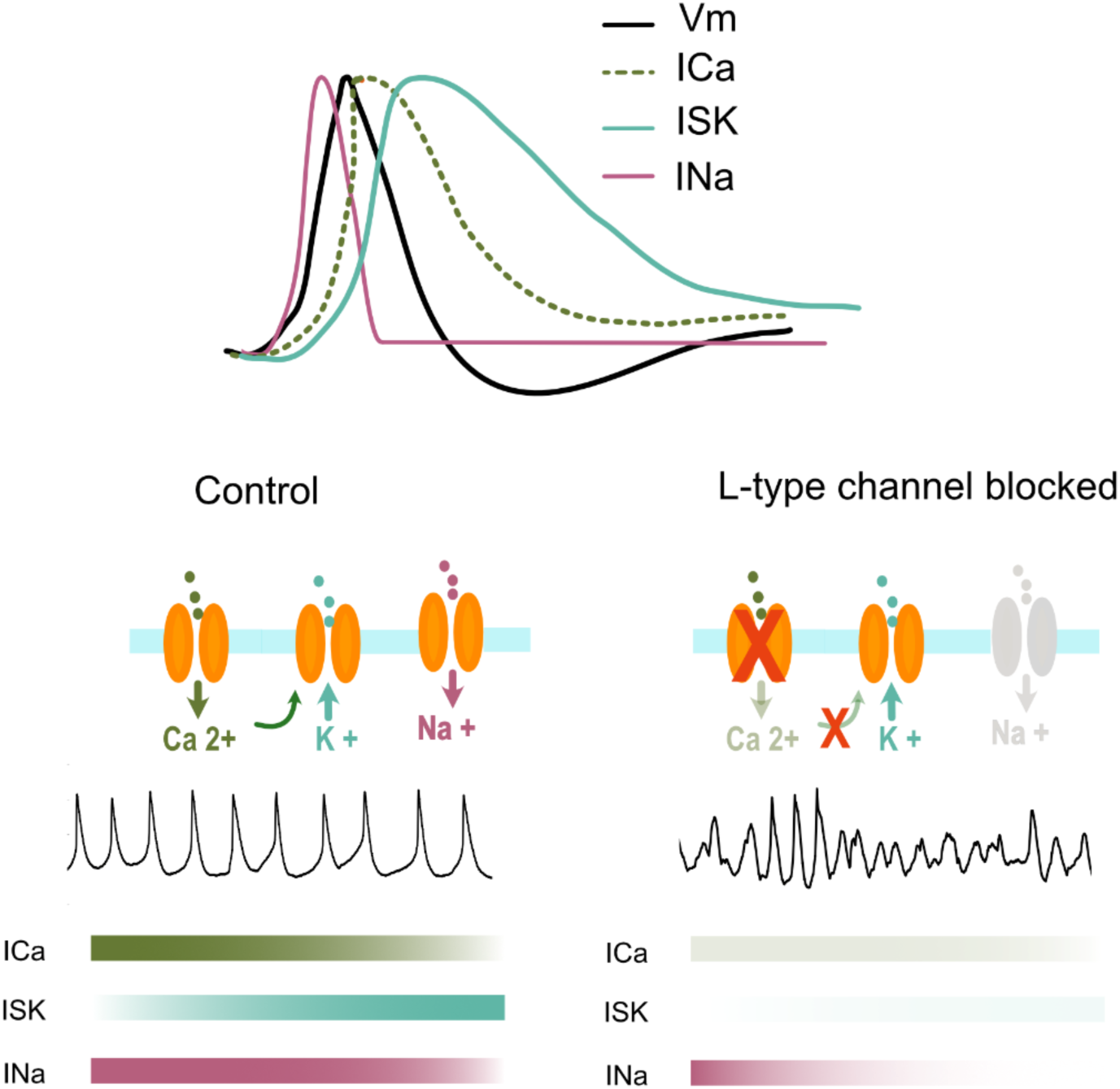

Legend: Top: The activation of voltage gated sodium channels (INa, pink), voltage gated calcium channels (ICa, dotted green), and SK-type channels (ISK, teal) during the action potential (Vm, black). Bottom: Under control conditions, sodium entry via voltage-gated sodium channels leads to depolarisation resulting in the activation of calcium channels. The elevation of intracellular calcium levels activates SK-type calcium-dependent potassium channels which repolarise the membrane, removing sodium channel inactivation. With L-type voltage-gated calcium channels blocked, this process is affected causing cessation of sustained tonic spiking.

## Introduction

The cerebellum is one of the most highly conserved regions of the vertebrate brain: the cell types, gene expression patterns and connectivity are highly homologous among all vertebrate classes from fish to mammals (Nieuwenhuys, 1967; Bell, 2002; Bae et al., 2009). This is particularly true for Purkinje neurons (PNs), which are the principal neurons of the cerebellar cortex and have similar molecular markers, dendritic morphology and connectivity across all vertebrates (Bae et al., 2009; Robra and Thirumalai, 2016; Kidd, 2017). PNs are critical for cerebellar function and their impairment leads to loss of motor co-ordination (Taroni and DiDonato, 2004; Chopra and Shakkottai, 2014). While the electrical properties of PNs have been well studied in mammals (Llinás and Sugimori, 1980; Raman and Bean, 1999; Womack and Khodakhah, 2003; Häusser et al., 2004), less is known in other vertebrates. Investigating the ionic basis of PN activity in non-mammalian vertebrates is important for understanding key properties essential for its functioning.

In zebrafish, PNs differentiate by 3 days post fertilization (dpf) (Bae et al., 2009; Hamling et al., 2015) and stable electrical activity patterns can be observed as early as 5 dpf (Sengupta and Thirumalai, 2015). In addition, calcium imaging and ablation studies report that the PN population participates in natural behaviours of larval zebrafish at these stages (Ahrens et al., 2012; Markov et al., 2021; Narayanan et al., 2024). Given the relative ease of cell identification, single cell recording and pharmacology in larval zebrafish, we chose to investigate the ionic basis of PN activity in this model system.

*In vivo* recordings show that like mammalian PNs, larval zebrafish PNs (zPNs) exhibit simple spikes mediated by TTX-sensitive sodium channels and that they can fire spontaneously even when isolated from the network (Sengupta and Thirumalai, 2015). Like mammalian PNs, these neurons also have elaborate dendritic arbors in the molecular layer and receive excitatory synaptic inputs from parallel fibers of granule cells and climbing fibers (CF) of the inferior olive (Bae et al., 2009; Sitaraman et al., 2021).

While the cerebellar circuitry and PN physiology are largely conserved across vertebrates (Fig. 1A), there are some differences between the firing properties of PNs in rodents versus zebrafish. Simple spikes are attenuated in the soma and complex spikes are completely absent, unlike mammalian PNs. Also, zPNs exhibit bistability, while this is a subject of debate for mammalian PNs (Loewenstein et al., 2005; Schonewille et al., 2006; Engbers et al., 2013; Sengupta and Thirumalai, 2015). These differences suggest underlying variation in the biophysics of PNs across these two systems. However, the kinds of ionic conductances expressed in zPNs and their contribution to spontaneous activity are largely unknown.

**Figure 1.**
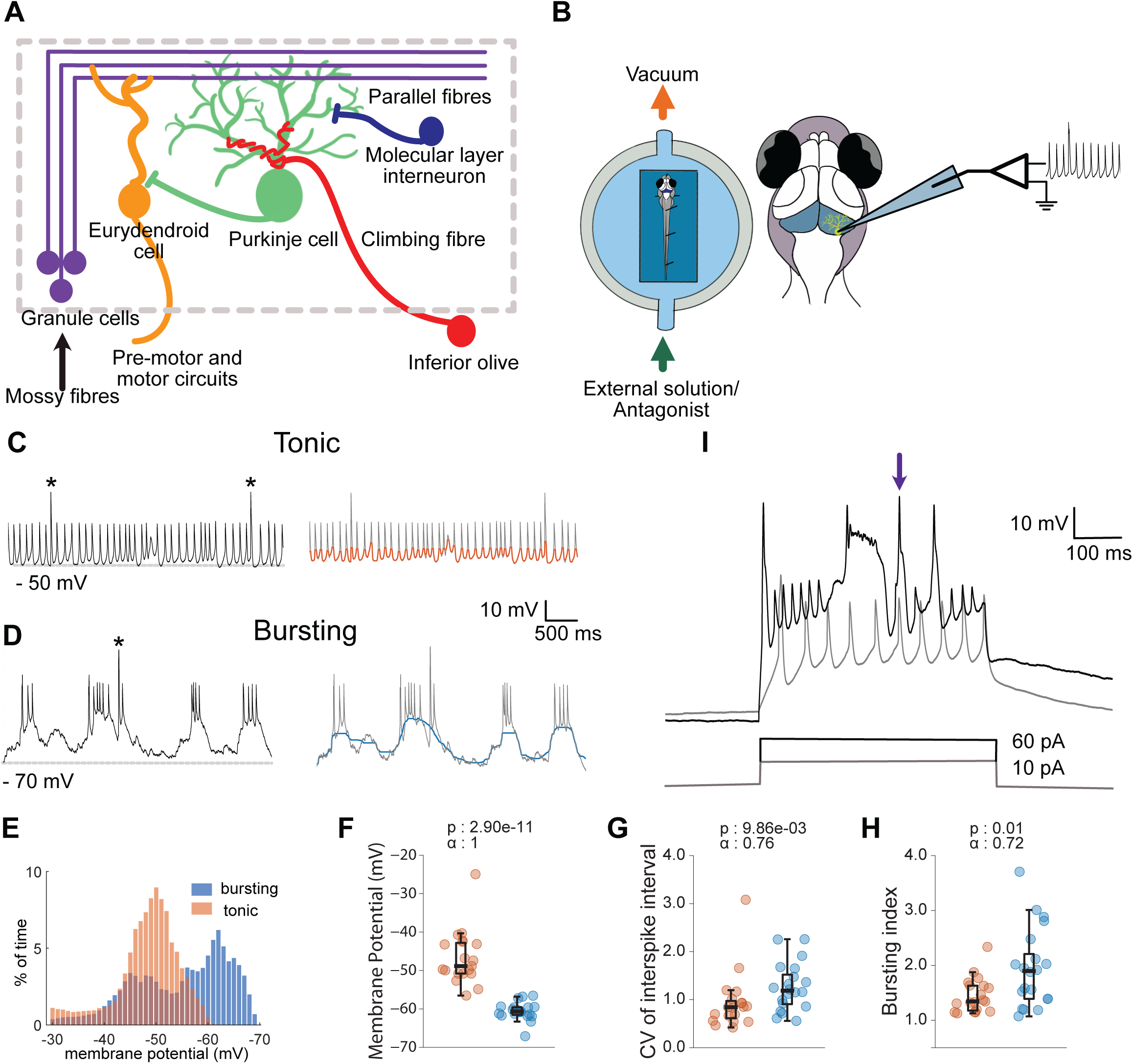
Spontaneous activity in Purkinje neurons. **A.** The cerebellar circuitry showing the major inputs to the PNs and their postsynaptic targets. **B.** Schematic showing the preparation for *in vivo* whole-cell recording of Purkinje neurons (PN). Location of the cerebellum in the larval zebrafish brain, highlighted in blue. **C.** Representative trace of a tonic firing cell (top). * Indicates climbing fibre driven EPSPs. The same trace after application of a moving median filter (window length 120 ms) (orange) to show membrane potential changes on a longer time scale (bottom). **D.** Same as C for a bursting cell. **E.** Distribution of membrane potentials of the traces in C and D after applying median filter. Tonic cell in orange and bursting in blue. **F.** Resting membrane potentials of cells in tonic and burst mode. For bursting, the lower membrane potential was used. **G.** Coefficient of variance of simple spike intervals in tonic and bursting cells. **H.** Bursting index for tonic and bursting cells (mean ISI/ median ISI), N=19 (tonic) and 20 (bursting), Mann-Whitney U test. **I.** Response of Purkinje neurons to depolarizing current pulses. Lower input current generates sodium driven simple spikes (gray trace) while a higher input current elicits large amplitude and broader calcium spikes (black trace). Arrow indicates a calcium spike.

Our earlier work showed that zPNs can exist in one of two stable membrane potential states: when in the depolarized state, they fire tonically and when hyperpolarized, they fire in bursts. Tonic firing is an intrinsic property of zPNs that persists even in the absence of synaptic inputs (Sengupta and Thirumalai, 2015), similar to mammalian PNs. Here, we investigate the ionic conductances responsible for tonic firing in zPNs. We show that these cells have a large calcium current contributed by L and T-type calcium channels with a minor contribution from P/Q type channels. Blocking L-type calcium channels or SK-type calcium dependent potassium channels had the same effect: a decrease or a complete cessation of tonic firing. Blocking SK channels also leads to a shallower AHP which reduces the pool of available sodium channels to drive tonic activity. Together, our results show that L-type and SK channels are critical for maintaining high-frequency tonic firing in zPNs.

## Results

### Spontaneous activity in Purkinje Neurons

We performed whole-cell patch clamp electrophysiology from PNs *in vivo* in larval zebrafish at 5-11 days post fertilization (dpf) (Fig. 1B). As has been reported previously, zPNs display two distinct firing modes: tonic and bursting (Fig. 1C, D). We were also able to switch the state of the cell by supplying a constant current. Both the modes show the presence of sodium-mediated simple spikes as well as climbing fiber (CF) EPSPs. The two modes also differ in several key characteristics: membrane potential, firing frequencies of simple spikes, and their dependence on synaptic input. We plotted the distribution of membrane potential at each data point from the traces in Fig. 1C and D, after applying a median filter to isolate the low frequency membrane potential changes. While the distribution for the tonic state is unimodal, that for the bursting state is bimodal (Fig. 1E). The modes of these distributions, referred to as resting membrane potential, are also different (Fig. 1F). The inter-spike intervals (ISI) of simple spikes are more variable for the bursting mode (Fig. 1G). This difference in inter-spike interval can also be seen as difference in the bursting index. Bursting index was quantified as the mean of ISIs divided by their median. Consequently, if the neuron is firing at regular intervals, the bursting index will be closer to 1. A higher bursting index indicates more variability in ISI and therefore more bursts (Fig. 1H).

Sengupta and Thirumalai, 2015 have shown that these cells can maintain tonic firing even in the absence of synaptic inputs. On the other hand, CF inputs can trigger bursts when zPNs are hyperpolarized. Further, zPNs display calcium spikes when they are depolarized above spike threshold (Fig. 1I, arrow). These results suggest that zPNs have significant high threshold calcium currents. It is also obvious that the tonic firing mode is dependent on the intrinsic properties of zPNs unlike the bursting mode which is activated by synaptic inputs.

Therefore, we wished to determine the major ion channels involved in maintaining tonic firing in these neurons. To do this, we measured cellular currents in voltage clamp mode and also compared spontaneous activity of zPNs when specific channel types were blocked with antagonists. We quantified several parameters including resting membrane potential, simple spike amplitudes, rates, shapes, etc., and compared them across the two conditions.

### Voltage-gated calcium channels regulate tonic firing

We wanted to understand how calcium currents shape the spontaneous activity of zPNs *in vivo*. For this, we first performed voltage clamp experiments to measure the total calcium current. As shown in Fig. 2A, we delivered voltage command pulses in the presence of TTX and TEA and measured cellular currents before and after application of 10 µM cadmium chloride, a non-specific, high voltage-gated calcium channel blocker. A cadmium-sensitive inward current with peak values at a holding potential of −20 mV was observed in all cells we recorded from (Fig. 2B, C).

**Figure 2.**
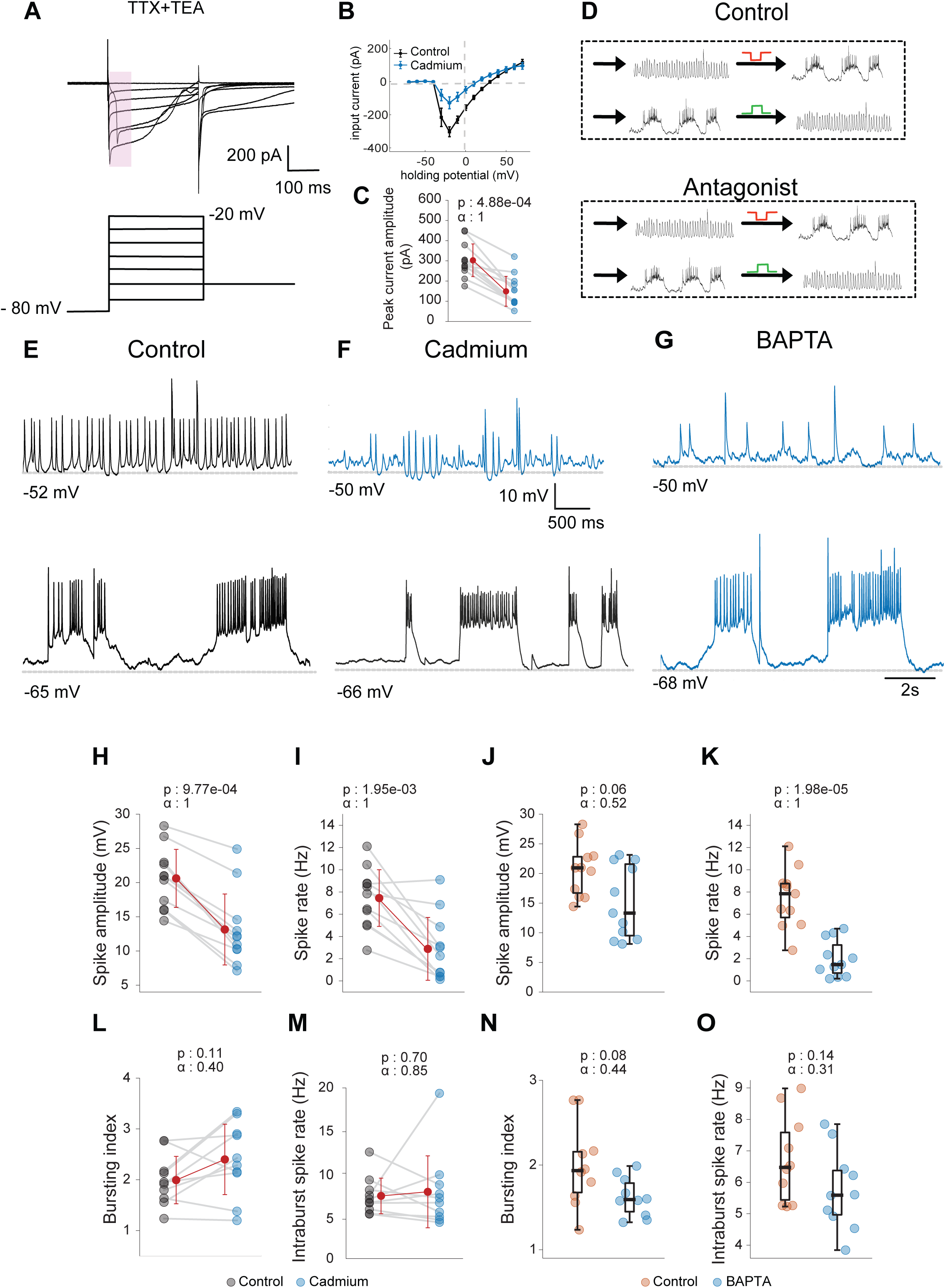
Blocking calcium channel affects tonic firing of simple spikes independent of synaptic inputs. **A.** Schematic representing the command voltage given and a corresponding representative current response of a cell in the presence of TTX (1 µM) and TEA bromide (1 mM). **B.** Average I-V response of cells before and after addition of cadmium chloride. **C.** The maximum inward current at −20 mV in cells before and after cadmium chloride (10 µM) N=12 cells, Wilcoxon signed rank test. **D.** Protocol used for paired spontaneous recordings of PNs. Each cell was recorded from in both tonic and bursting state before (control) and after addition of the channel antagonist. **E and F.** Representative traces of tonic and bursting cells before (black) and after addition of cadmium chloride (10 µM) (blue). **G.** Representative traces of cells firing in tonic and in burst mode with BAPTA in the recording pipette. **H-I.** Mean simple spike amplitude and spike rate in tonic firing cells without and with cadmium., N = 11 cells, Wilcoxon signed rank test. **J-K.** Mean simple spike amplitude and spike rate in tonic firing cells in control cells versus BAPTA. N = 11 cells, Mann-Whitney U test. **L-M.** Bursting index and intra-burst simple spike rate before and after cadmium chloride. N=10 cells, Wilcoxon signed rank test. **N-O.** Bursting index and intra-burst simple spike rate in control and BAPTA. N=10 cells, Mann-Whitney U Test.

Next, we recorded from zPNs in current clamp mode under bursting and tonic-firing conditions before and after application of 10 µM cadmium chloride. We recorded the state of the cell without current injection and then made it switch states by injecting a constant depolarizing or hyperpolarizing current. After perfusing the antagonist in the bath, we recorded from the same cell again in both states (Fig. 2D).

We found that the cells could still fire in both states even in the presence of cadmium (Fig. 2E, F). 4 out of 11 cells switched states upon the addition of cadmium while the rest continued to fire in the same state. Although the resting membrane potential in both states did not change, tonic firing of simple spikes was affected. Simple spikes became smaller in amplitude and less frequent (Fig. 2H, I). ISIs became more irregular, with a higher coefficient of variation: CV of ISI in tonic mode was 0.81 ± 0.06 (control) and 1.73 ± 0.11 (cadmium); (Mean ± SD). 3 of the 11 cells went into depolarization-induced block later in the recording. We did not observe any drastic changes in the bursting behavior and the bursting index remained the same in cadmium compared to control (Fig. 2L, M).

The above results suggest a role for voltage-gated calcium channels in tonic firing. However, application of cadmium in the bath affects voltage-gated channels everywhere, including those that mediate neurotransmission. To rule out network-mediated effects and to test this in a cell specific manner, we next blocked calcium channels intracellularly using the calcium chelator 1,2-bis (o-amino phenoxy) ethane-N, N, N′, N′-tetra acetic acid (BAPTA). We recorded from cells using normal patch internal solution or with 20 mM of BAPTA. Cells recorded with BAPTA in internal solution were also able to fire in both states (Fig. 2G), with a decrease in frequency of tonic simple spikes (Fig. 2K). Spike amplitude in BAPTA was not altered with a borderline p-value of 0.06 (Fig. 2J). Bursting index and the spike rate within bursts remained unaffected (Fig. 2N, O), underlining the importance of calcium currents in regulating tonic firing specifically.

### Calcium currents in zPNs are primarily driven by L and T-type channels

Neurons express many different types of voltage-gated calcium channels, with distinct kinetics and effects on neuronal firing properties. We wanted to determine the classes of voltage-gated calcium channels expressed in zPNs. Takeuchi et al., 2017 analyzed the transcriptome of 14 dpf zPNs. From their dataset, we looked for the expression levels of genes encoding voltage-gated calcium channels and found significant expression of transcripts encoding L-type, P/Q type and T-type channel subunits, with L-type showing highest levels of expression (Fig. 3A). Auxiliary subunits of calcium channels were also strongly expressed.

**Figure 3.**
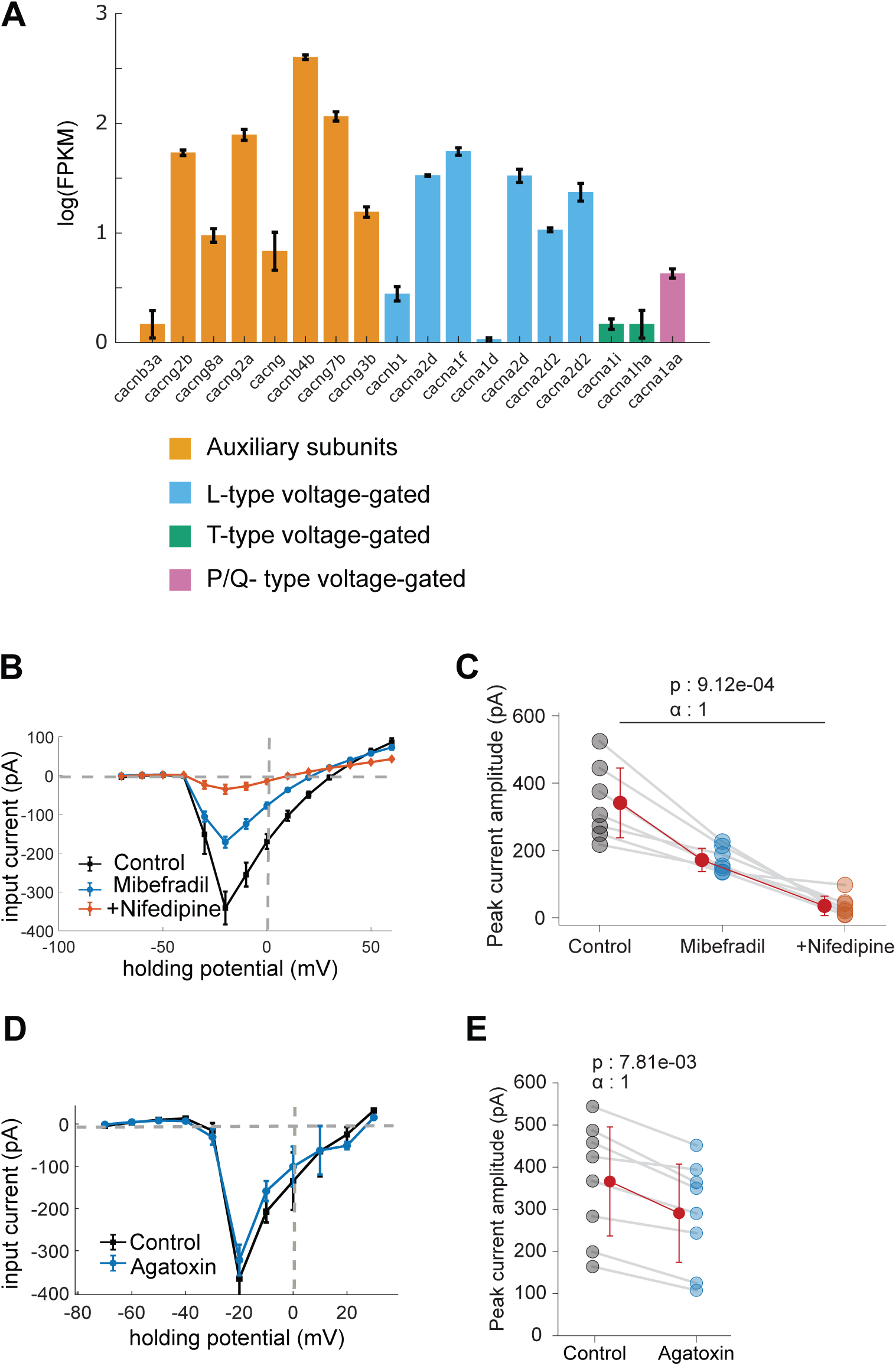
Calcium current in Purkinje cells is mainly contributed by L and T type voltage gated calcium channels. **A.** mRNA levels of genes for voltage-gated calcium channels in 12 dpf larval zebrafish. (Data from Takeuchi et. al., 2017). **B and D.** Average I-V response of cells. All recordings were done in the presence of TTX and TEA. Without antagonist (black), with T-type channel antagonist mibefradil (blue) (100 µM) followed by L-type channel antagonist, nifedipine (orange) (200 µM). (B). Before and after addition of ω-agatoxin IVA (100 nM) (D). **C and E.** The maximum inward current at −20 mV in cells in control, with mibefradil and nifedipine, N=7 cells, Friedman’s test (C). Maximum inward current before and after addition of ω-agatoxin IVA, N=8 cells, Wilcoxon signed-rank test (E).

Supported by this dataset, we set out to measure calcium currents in zPNs using voltage clamp protocols and pharmacology. We recorded the current response of cells to a series of voltage pulses in the presence of 1 µM TTX and 1 mM TEA. We recorded the response in normal saline and after application of different calcium channel antagonists and plotted their current-voltage (I-V) relationships. As before, we observed a large inward current active between −50 mV and +20 mV (Fig. 3B and D). We then quantified the contributions of three subtypes-L-, T- and P/Q-types using nifedipine, mibefradil and ω-agatoxin IVA, respectively. Nifedipine has been shown to have off target effects on T-type channels (Shcheglovitov et al., 2005). To nullify the off-target effect, we first added mibefradil, a T-type specific blocker (Esneu et al., 1998), followed by nifedipine. We found that the inward current was partially blocked by the T-type channel blocker, mibefradil. In 100 µM mibefradil, the peak inward current was reduced to 50% of the control values. Addition of 100 µM of the L-type channel blocker nifedipine further decreased the peak inward current to 10% of the control values (Fig. 3B, C). On the other hand, application of the P/Q type channel blocker, ω-agatoxin IVA (100 nM) reduced the current by only 9 percent (Fig. 3D, E). We also found that the presence of ω-agatoxin IVA had no significant effect on the tonic firing rate in Purkinje neurons (control: 7.78 ± 1.75 Hz; ω-agatoxin IVA: 6.90; ± 1.68 Hz; Mean ± SD, N= 6; data not shown).

Overall, we found that L and T-type calcium channels are major contributors to voltage-gated calcium current. Wen and group (Wen et. al., 2013) have reported a reduced sensitivity of ω-agatoxin IVA to zebrafish P/Q channels. It is therefore possible that we underestimated the P/Q current in our voltage-clamp studies. However, both voltage-clamp and transcriptome data show a major contribution of L-type calcium channels.

We next asked how each of these channel types contributed to the tonic firing in these cells.

### L-type, but not T-type channels are required for sustained tonic firing

To understand the contribution of T-type calcium channels to tonic firing in zPNs, we recorded their firing pattern before and after application of the T-type channel blocker, mibefradil. Application of mibefradil reduced spike amplitude but had no effect on spike frequency (Fig. 4B, C).

**Figure 4.**
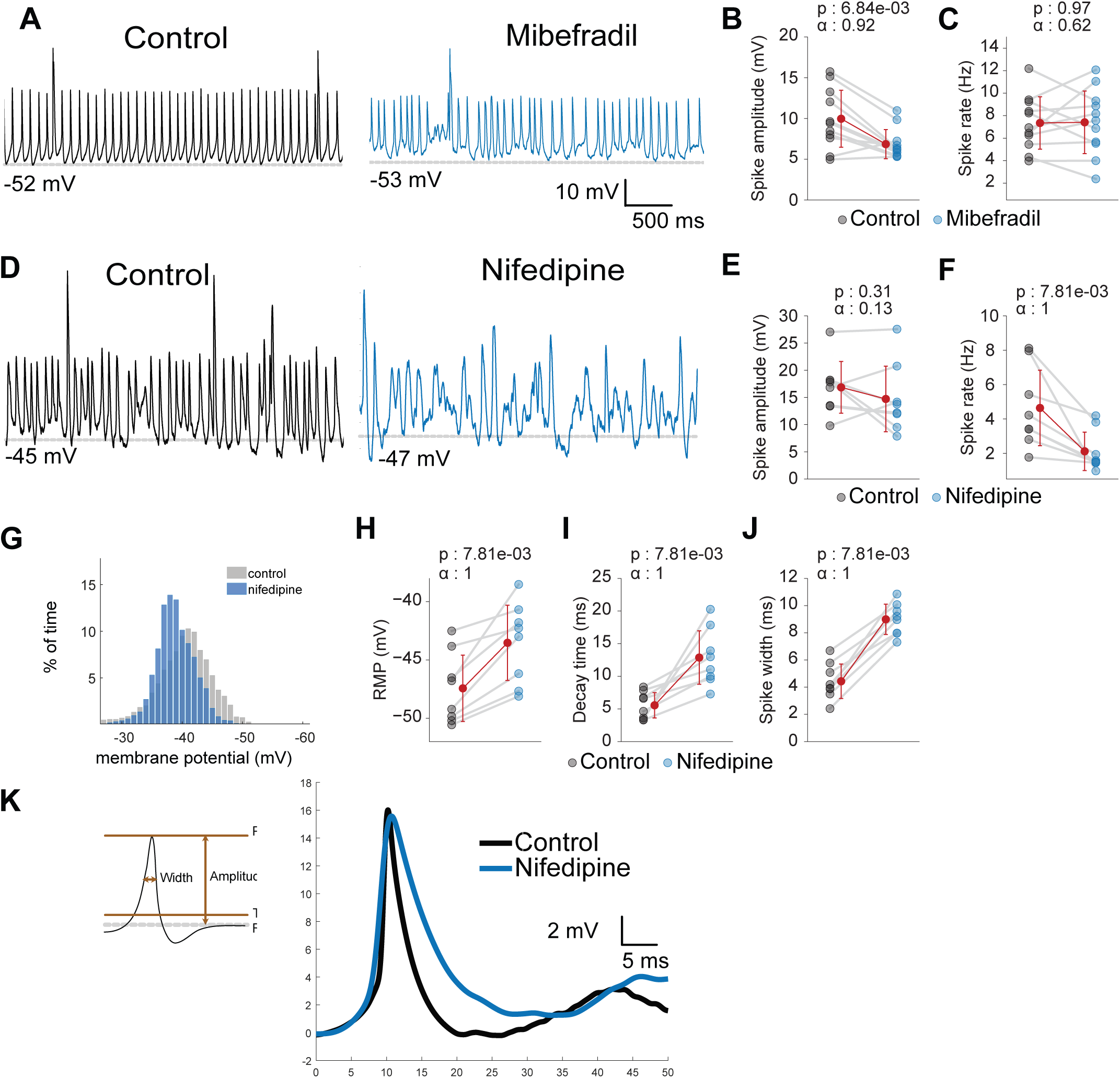
L-type and not T-type channel are important for tonic firing and membrane repolarization. **A.** Representative traces of tonic firing before and after addition of T-type antagonist, mibefradil (100 µM). **B-C.** Mean simple spike amplitude and spike rate in control and in presence of mibefradil. N= 11; Wilcoxon signed-rank test. **D.** Representative traces of tonic firing before and after addition of L-type antagonist, nifedipine (200 µM). **E-F.** Mean, simple spike amplitude and spike rate in control and in the presence of nifedipine N= 8; Wilcoxon signed-rank test. **G.** Distribution of membrane potential for one representative cell before and after addition of nifedipine. **H.** Mean resting membrane potential in control and in the presence of nifedipine. **I-J.** Mean decay time and width of simple spikes during tonic firing before and after addition of nifedipine. N= 8; Wilcoxon signed-rank test. **K.** Averaged simple spike waveform from all simple spikes in one representative cell before (black) and after addition of nifedipine (blue).

Next, we recorded from zPNs in current clamp mode before and after application of the L-type channel blocker, nifedipine. The effect on spike rate was similar to that of adding cadmium (Fig. 4D-F). Nifedipine also changed the shape of the spike, as evidenced by increases in spike decay time and width (Fig. 4I-K). To determine if this was due to overall changes in membrane potential, we looked at the distribution of membrane potentials in control versus presence of nifedipine. We observed an overall shift towards more depolarized membrane potentials across all cells (Fig. 4G) with a significant increase observed in the resting membrane potential (RMP, Fig. 4H). These results are consistent with the broadening of the simple spike and the overall reduction in spike rate.

### L-type calcium currents are active during spike repolarization

To better understand the effect of nifedipine without the influence of synaptic inputs, we also recorded the response of cells in the presence of synaptic blockers (NBQX and gabazine). We then provided a negative current to hyperpolarize the cell to −70 mV. In this way, the cells were silent unless depolarized (Fig. 5A). We observed that the amplitude of AHP decreased while the threshold to spiking increased further confirming the role of L-type channels in repolarization (Fig 5C-D). In response to a 2-second-long depolarizing current, there was an increase in the spike adaptation ratio (Frequency of first spike/ Frequency of 10th spike) further suggesting that this repolarization maybe important for sustained firing of simple spikes (Fig. 5A, E).

**Figure 5.**
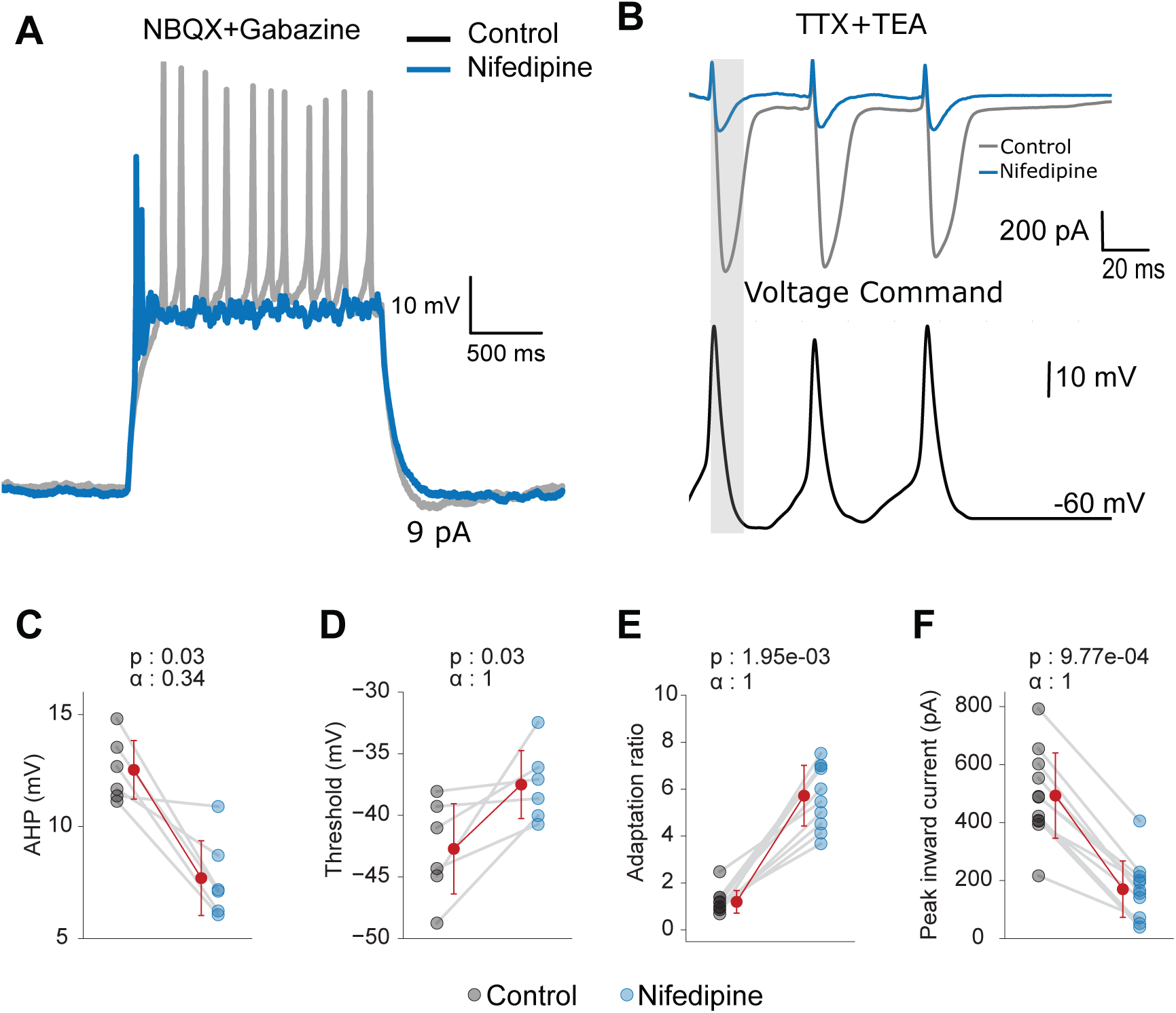
L-type channels are active during the repolarization phase of simple spikes **A.** Representative response of a cell to a depolarizing pulse of current before and after addition of nifedipine (200 µM). These experiments were done in the presence of synaptic blockers NBQX (20 µM) and Gabazine (10 µM). A constant negative current was applied to keep the cell membrane potential at ∼-70 mV. **B.** Simple spike waveform given as voltage command and the representative current response of a cell without and with nifedipine. Experiments were performed in the presence of TTX (1 µM) and TEA (1 mM). **C-E.** Mean AHP amplitude, spike threshold and adaptation ratio before and after addition of nifedipine. N= 6; Wilcoxon signed-rank test. **F.** Peak inward current in response to simple waveform as a voltage command without and with nifedipine. N= 10; Wilcoxon signed-rank test.

Calcium imaging studies from PNs show calcium entry and elevation of cytosolic calcium levels during simple spike firing (Ramirez and Stell, 2016, Varma et al., 2024, Knogler et al., 2019), suggesting that voltage-gated calcium channels are activated during simple spike firing. To confirm if L-type channels are indeed active during simple spike firing in zPNs and to explore their possible role in repolarization, we replayed a train of simple spikes as the voltage command. We found that replaying the simple spike waveform as recorded at the soma (e.g., Fig. 1C) did not recruit sodium channels. This could be because of attenuation of the somatically recorded spike waveform, making the waveform too weak to activate sodium channels in the axonal spike initiation zone. Scaling up the amplitude of simple spikes led to a significant sodium current (data not shown). We, therefore, decided to use a scaled-up version of the simple spike waveform (∼2.5 x of original).

To quantify the calcium currents, all experiments were done in the presence of sodium and potassium channel antagonists (TTX and TEA). We found a large, inward current in response to the simple spike waveform. We also saw that this current was slow to activate and achieved maximum amplitude well beyond the peak of a simple spike, corresponding to the repolarization and AHP phases of the simple spike (Fig. 5B). Importantly, this current decreased by 66% in the presence of nifedipine (Fig. 5B and F).

These experiments show that L-type channels are indeed active when the cell fires simple spikes. The channels achieve maximum activation after the peak of the simple spike, during the repolarization phase.

### BK channels also contribute to membrane repolarization but do not modulate tonic firing frequency

We were surprised to find that the L-type calcium channels were active during the repolarization phase and that blocking them increased spike width. Both of these observations could be explained if these effects were mediated by calcium-dependent potassium channels (KCa). If these were true, the effect of blocking KCa channels should mimic the effect of blocking L-type calcium channels. Typically, KCa channels can be of large (BK), intermediate (IK) or small (SK) conductance. Of these BK and SK channels are known to be important for spike repolarization and AHP (Womack and Khodakhah, 2003; Edgerton and Reinhart, 2003).

We first tested whether BK channels couple with L-type channels and contribute to membrane repolarization. Blocking BK channels with 100 nM iberiotoxin, had no effect on the tonic firing frequency of these cells (Fig. 6A, C). The spike amplitude was also not affected (Fig. 6B). However, similar to L-type block, we observed a slight broadening of the simple spike and a dip in the AHP amplitude (Fig. 6D-G).

**Figure 6.**
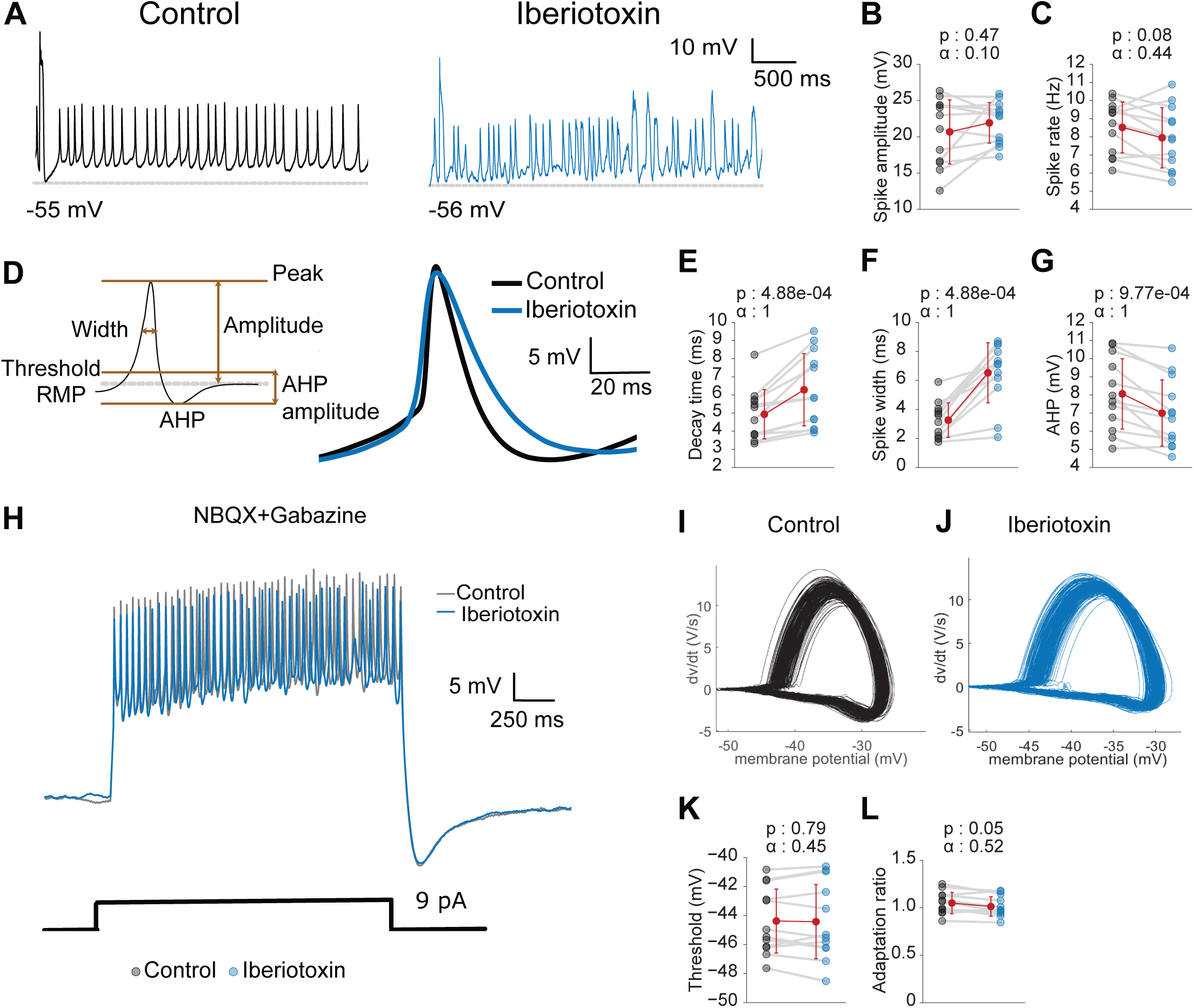
BK channels also contribute to membrane repolarization but do not modulate the tonic firing frequency. **A.** Representative trace of a tonic firing cell before and after addition of the BK channel antagonist, iberiotoxin (100 nM). **B-C.** Mean simple spike amplitude and spike rate before and after addition of iberiotoxin. N=12; Wilcoxon signed-rank test. **D.** Averaged simple spike waveform from one representative cell before (black) and after addition of iberiotoxin (blue). **E-G.** Mean decay time, width and AHP amplitude of simple spikes before and after addition of iberiotoxin. N=12; Wilcoxon signed-rank test. **H.** Representative response of a cell to a depolarizing pulse of current before and after addition of iberiotoxin. These experiments were done in the presence of synaptic blockers NBQX (20 µM) and Gabazine (10 µM). A constant negative current was applied to keep the cell membrane potential at ∼-70 mV. **I-J.** Phase plane of all simple spikes in a representative cell in control and with iberiotoxin. **K-L.** Mean spike adaptation ratio and spiking threshold in control condition and with iberiotoxin. N=12; Wilcoxon signed-rank test.

We also examined whether zPNs could generate simple spikes in a sustained manner upon depolarization, before and after application of iberiotoxin (Fig. 6H). These experiments were performed in the presence of synaptic blockers and the cell was depolarized from a baseline of −70 mV. Presence of iberiotoxin did not change the spike adaptation ratio (Frequency of first spike/ Frequency of 10th spike) or spiking threshold (Fig. 6I-L). This suggests that while BK channels may contribute to membrane repolarization, they do not control the firing frequency of simple spikes.

### SK channels couple to L-type channels to sustain tonic firing

On the other hand, we observed that blocking small conductance (SK) channels with apamin mimicked the effects of L-type channel block (Fig. 7A). Simple spikes were smaller and less frequent in the tonic mode (Fig. 7 D, E) despite only a small increase in the resting membrane potential (Fig. 7C). The membrane potential distribution also revealed further similarities between the apamin and nifedipine conditions (Fig. 7B). Similar to experiments with nifedipine, we observed a shift in the membrane potential as compared to control with fewer points in spiking range (−40 to −35 mV). This indicated that while the cells were slightly depolarized in the presence of apamin they did fire simple spikes less frequently. Spikes also became broader and the AHP shallower pointing toward their role in the repolarization of simple spikes (Fig. 7 F-I).

**Figure 7.**
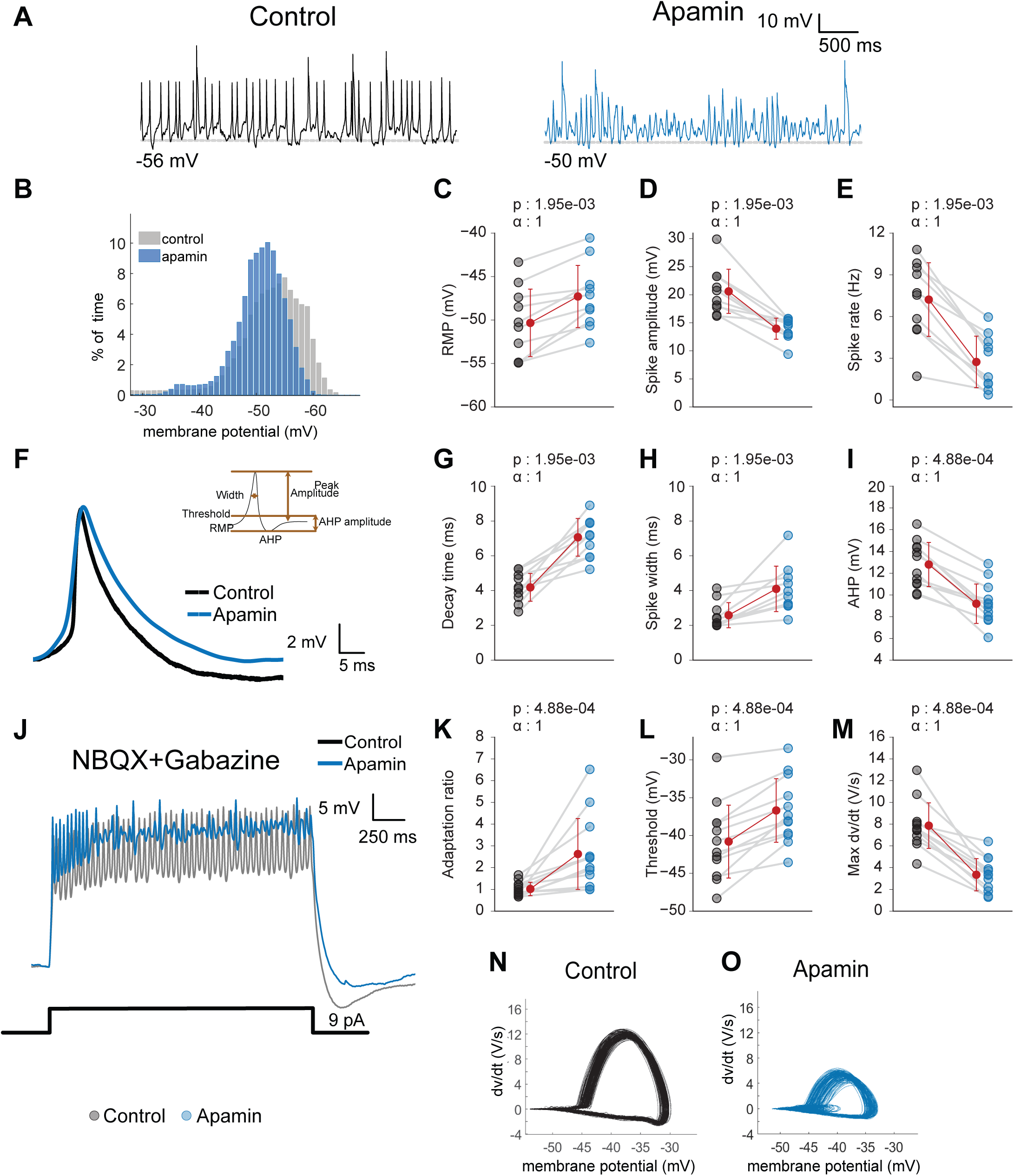
SK channels allow cells to maintain tonic firing by regulating sodium channel availability. **A.** Representative trace of a tonic firing cell before and after addition of the SK channel antagonist, apamin (200 nM). **B.** Distribution of membrane potential of a representative cell before and after addition of apamin. **C-E.** Mean resting membrane potential, simple spike amplitude and spike rate before and after addition of apamin. N=10; Wilcoxon signed-rank test. **F.** Averaged simple spike waveform from one representative cell before (black) and after addition of apamin (blue). **G-I.** Mean decay time, width and AHP amplitude of simple spikes. N=10; Wilcoxon signed-rank test. **J.** Representative response of a cell to a depolarizing pulse of current before and after addition of apamin. These experiments were done in the presence of synaptic blockers NBQX (20 µM) and Gabazine (10 µM). **K-M.** Mean spike adaptation ratio, spiking threshold and maximum dv/dt of simple spikes in response to current pulse in control condition and with apamin. **N-O.** Phase plane of all simple spikes in a representative cell in control and with apamin. N=10; Wilcoxon signed-rank test.

These data show that L-type calcium channels maintain the tonic firing of simple spikes and contribute to spike repolarization by activating SK channels.

### SK currents sustain high-frequency firing for a long duration by increasing sodium channel availability

The input resistance of the cells did not change significantly in the presence of either apamin or nifedipine (control: 1.559 ± 0.134 GΩ; apamin: 1.078 ± 0.154 GΩ; Mean ± SD; p-value: 0.912). There was no change in rheobase either (control: 4.600 ± 2.30 pA; apamin: 4.800 ± 1.14 pA; Mean ± SD; p-value 0.2266; Wilcoxon signed-rank test; N= 10). To test if SK channels contribute to sustained firing, we depolarized zPNs and recorded their firing responses in the absence of synaptic input as described in the previous section. In response to a two-second-long pulse, the cells did not show any spike adaptation across the entire pulse under control conditions. On the other hand, in the presence of apamin, the same cells fired simple spikes for up to 500 ms and then became irregular or silent (Fig. 7J). On average, the cells fired fewer spikes in apamin and had a higher spike adaptation ratio (Fig. 7K).

Lower repolarization and shallower AHP can reduce sodium channel recovery from inactivation. With time, this can reduce the availability of these channels and prevent the cell from generating more spikes. This usually leads to a depolarized threshold and a decrease in the maximum dv/dt of a spike (Colbert et al., 1997; Vandael et al., 2012). We did indeed observe this with apamin (Fig. 7L-O).

Overall, these data show that L-type calcium channels and SK calcium-dependent potassium channels are important for sustaining tonic firing for longer durations. They do so by allowing membrane repolarization and maintaining sodium channel availability.

## Discussion

The larval zebrafish provides an excellent system where neuronal physiology can be studied *in vivo*, in an intact circuit. Our study sheds some light on how ionic mechanisms drive tonic firing in an *in vivo* preparation. Here, we have shown that zPNs express interesting membrane potential dynamics driven by diverse channel types. We identified high- and low-voltage-gated calcium currents and calcium-activated potassium currents in these neurons. Together with SK-type KCa channels, L-type calcium channels contribute to the repolarization of membrane potential and AHP of simple spikes. They allow cells to fire tonically for a long duration in the up state by increasing the availability of sodium channels. Overall, we show that L-type voltage-gated calcium channels and SK calcium-activated potassium channels are important in maintaining tonic firing in the up state of zPNs. Further work can help shed light on the ionic mechanisms of bistability in larval zebrafish Purkinje neurons.

### Limitations of the study

Pharmacological studies are most limited by a lack of specificity. We applied each antagonist in the external milieu of the fish, blocking a class of channels throughout the body. Because of this, when comparing firing patterns, we cannot completely negate the effect of altered synaptic inputs. Nevertheless, our BAPTA experiments show that the effects of calcium channel block on simple spike firing rate are mainly due to intrinsic changes. Further, the effects of nifedipine and apamin on tonic firing persisted even under conditions of synaptic blockade. These results give us confidence that we have successfully isolated ionic mechanisms of tonic firing in zPNs.

Still, we cannot rule out any off-target effects of the channel antagonists or that they have reduced affinity for their intended targets in zebrafish. To address the latter, we performed voltage clamp experiments to test if the blockers affected cellular currents in their expected voltage range. For example, the L-type channel blocker nifedipine blocked cellular currents in the depolarized voltage ranges (Fig. 3B), as expected. These results attest to the specificity of these agents towards their intended target. The relatively minor effects of ω-agatoxin IVA and iberiotoxin could also be due to their limited interactions with zebrafish P/Q-type and BK channels respectively and therefore, these negative results should be interpreted cautiously. Further, even in mammalian PNs, the BK current is known to comprise an iberiotoxin-insensitive component (Benton et al., 2013), mediated by the β4 subunit (Meera et al., 2000). This raises questions whether this is the case in zPNs as well. However, a recent study showed that the β4 subunit gene *kcnmb4* is lost in zebrafish (Silic et al., 2021). Additionally, transcriptome data from Takeuchi et. al., 2017 indicates significant expression of only iberiotoxin-sensitive BK1 channels, suggesting that in zPNs, an iberiotoxin-insensitive BK current may be absent. This study has also not examined persistent sodium channels, which are known to drive tonic firing in other neurons (Agrawal et. al., 2001; Cho et. al., 2015), including mammalian PNs (Raman and Bean, 1997).

In this study, we focused on channels that affected the tonic state. None of the identified channels seemed to affect spiking in the bursting state. Bursting requires synaptic input and when GABAergic and glutamatergic inputs are blocked, zPNs become quiescent upon hyperpolarization. A full investigation of mechanisms driving bursting and those governing switches from one state to another is necessary to fully understand PN dynamics. One challenge is the high variability of bursting within and among cells. A higher sample size (both in the duration of recordings and the number of cells) might allow us to find subtle changes in bursting parameters.

Our study captures the contribution of calcium currents in a small developmental window. All of the experiments were done on larvae between the ages of 5-11 dpf. By this stage, the cerebellum is functional and Purkinje neurons display their characteristic firing. However, this is still an early developmental stage and the circuit undergoes further maturation. The cerebellum grows in volume with a concurrent increase in Purkinje neuron numbers (Bae et al., 2009; Hamling et al., 2015). During this time, PNs elaborate dendritic arbors in the molecular layer and make numerous synaptic contacts with parallel and climbing fibers (Sitaraman et al., 2021). But not much is known about maturation of ion channel distributions in zPNs. In rodents, it is well appreciated that the first two weeks of post-natal life is a period of drastic developmental changes in the cerebellum (van Welie et al., 2011) with changes in firing properties (McKay and Turner, 2005), ion channel distributions (Womack and Khodakhah, 2003; Hosy et al., 2011), and synaptic pruning (Hashimoto and Kano, 2013). It is likely that zPNs also undergo such developmental transformation during early larval stages and these processes remain to be studied.

### Comparative physiology of Purkinje neurons in teleosts and mammals

Purkinje neurons in teleosts share many of the properties of mammalian PNs as has been shown in this and in many previous studies. PNs in teleosts and in mammals are arranged in a monolayer and send elaborate dendritic arbors into the molecular layer where they receive numerous excitatory and inhibitory synaptic inputs from conserved cell types and pathways (Meek, 1992). Studies localizing protein markers have been carried out extensively in rodent PNs and in zPNs. Both rodent PNs and zPNs express the dopamine and cAMP regulated phospho protein Darpp32 also known as ppp1r1b (Ouimet et al., 1984, 1992; Robra and Thirumalai, 2016). Expression of Glutamate receptor delta 2 is also seen in both cases (Araki et al., 1993; Mikami et al., 2004; Tam et al., 2021). In contrast, Zebrin II expression shows distinct expression patterns in rodent and zebrafish PNs: in rodents, PNs are organized into alternating Zebrin II -positive and negative stripes whereas in zebrafish, the expression is uniform in all PNs (Brochu et al., 1990; Bae et al., 2009).

Both mammalian PNs and teleost PNs receive excitatory inputs from parallel fibers and climbing fibers and fire simple spikes spontaneously (Sengupta and Thirumalai, 2015; Pose-Méndez et al., 2023). In rodents, PNs seem to exhibit bistable behavior, with an ‘up’ state where they fire action potentials and a ‘down’ state where they are quiescent and may fire bursts (Williams et al., 2002; Womack and Khodakhah, 2002). Excitatory climbing fiber inputs or inhibitory input from molecular layer interneurons could toggle PNs between these states (Loewenstein et al., 2005; Oldfield et al., 2010; Engbers et al., 2013). Nevertheless, bistability in rodents PNs *in vivo* is contested due to the effect of anesthetics on PN physiology (Schonewille et al., 2006). We have shown previously that zPNs are bistable and that stimulating climbing fibers triggers bursts (Sengupta and Thirumalai, 2015).

Finally, it is noteworthy that zebrafish are poikilotherms and prefer to live at temperatures around 25 to 30°C. Under lab conditions, they are maintained at 28 °C, which is much cooler than the mammalian body temperature of 37 °C. Such differences in temperature might contribute to large differences in channel kinetics. Despite this, activity patterns not only in PNs but also other cell types such as spinal motor neurons are largely similar across zebrafish and mammals. For instance the intracellularly recorded spike width of cat motor neurons is comparable to that of larval zebrafish motor neurons (Brock et al., 1952; Jha and Thirumalai, 2020). We speculate that compensatory mechanisms could exist to mitigate the temperature differences. These could occur at two levels: the structure of the channel could be modified such that the gating kinetics remain consistent despite the difference in temperature; or the difference in kinetics of channels may be compensated such that overall output of the neuron remains the same. Indeed, studies have shown that cold-blooded systems are capable of maintaining similar output of neurons or networks over a wide range of temperatures (Alonso and Marder, 2020).

### Calcium and calcium-activated potassium currents in PNs

Much of what we know regarding ionic conductances underlying PN activity comes from experiments performed in rodent cerebellar slices acutely, or PNs in dissociated culture. These studies have shown that calcium currents activated by the action potential are mainly carried by P/Q-type and T-type voltage-gated calcium channels (Raman and Bean, 1999; Swensen and Bean, 2003). Calcium entry through these channels activates BK and SK calcium-dependent potassium channels and for this reason, blocking calcium channels or KCa potassium channels produces the same effect on spontaneous firing of PNs (Womack and Khodakhah, 2002, 2003; Edgerton and Reinhart, 2003). Our experiments on zPNs point to some similarities and differences. Like in mammals, fish PNs also regulate spontaneous tonic firing via calcium channels and calcium-dependent potassium channels. But, in zPNs, calcium enters via the L-type and not the P/Q type calcium channels. While P/Q type channels are coupled to SK channels in rodents, we find that L-type channels are coupled to SK channels in fish. Neuronal firing is regulated by the coupling of calcium channels and calcium-dependent channels in many other neuronal types as well such as in hippocampal pyramidal neurons (Lancaster et al., 1991), midbrain dopaminergic neurons (Wolfart and Roeper, 2002), neurons in the medial vestibular nucleus (Smith et al., 2002) etc. In each case, the voltage-gated calcium channel through which calcium enters and the type of calcium-dependent potassium channels it is coupled to is different. Together with our results, these studies suggest that coupling of calcium and calcium-dependent potassium channels is a conserved mechanism for regulating neuronal firing properties across vertebrates and across neuronal types.

## Methods

### Ethical Approval

The use of all experimental animals (*Danio rerio*) was approved by the Institutional Animal Ethics Committee (NCBS-IAE-2020/14(R1E)) and the Institutional Biosafety Committee (TFR/NCBS/14-IBSC/VT1/2011).

### Fish rearing and care

Indian wild type zebrafish were used for all experiments. These were originally sourced from local aquariums and then the generations were propagated and maintained in-house. All adults were housed in a continuous recirculating system (ZebTec, Tecniplast, Italy). Conductivity and pH of water were maintained at 1200 μS and 7.2 respectively. The adults were bred to obtain embryos. Once collected, embryos were maintained in small tanks without constant circulation. Methylene blue (0.0005%) was added to the water from the recirculation system and this was used to raise the larvae until the day of the experiments. Adults were fed Zeigler diet and brine shrimp larvae while larvae were fed Zeigler Larval Diet. Both adults and larvae were maintained at 28 °C with a 14:10 hour light: dark cycle. All experiments were performed on larvae at 5-11 dpf. After completion of the experiment, the larvae were euthanized with rapid chilling.

### Whole-cell patch clamp

#### Animal preparation

Larvae were anesthetized with 0.015% MS222 (tricaine) and then transferred onto a block of Sylgard (Dow Corning, Midland, MI, United States) in the middle of the recording chamber. Tungsten wire (California Fine Wire, Grover Beach, CA, United States) was used to pin the tail, head and jaws to the Sylgard block. Next, the MS222 was replaced with external saline (composition in mM: 134 sodium chloride, 2.9 potassium chloride, 1.2 magnesium chloride, 10 HEPES, 10 Glucose, 2.1 calcium chloride, 0.01 alpha tubocurarine; pH: 7.8; 290 mOsm). Using a pair of fine forceps, the skin covering the brain was gently peeled off.

#### Electrodes and cell identification

Pipettes were pulled from thick-walled borosilicate capillaries (1.5 mm OD; 0.86 mm ID; Harvard Apparatus, Massachusetts, United States) using a Flaming Brown P-97 pipette puller (Sutter Instruments, Novato, CA, United States) such that they had a tip diameter of ∼1 µm and resistances in the range of 12-17 MΩ when filled with internal solution. Large, elliptical cells in the dorsal region of the cerebellum were targeted for whole-cell patch clamp. They were confirmed as Purkinje neurons based on the large CF-EPSCs and their characteristic firing profile in current clamp (Sengupta and Thirumalai, 2015). The recording solution contained sulphorhodamine (5 µg/mL) and the fill of the cell was also observed to confirm the characteristic spiny dendritic morphology of Purkinje neurons.

#### Recording solutions

For current clamp recordings, pipettes were filled with potassium gluconate-based internal solution (composition in mM: 115 potassium gluconate, 15 potassium chloride, 2 magnesium chloride, 10 HEPES, 10 EGTA, 4 Mg-ATP; pH: 7.2; 290 mOsm). In experiments where 20 mM tetra K-BAPTA (Life Technologies) was added, concentration of potassium gluconate was reduced to 35 mM with the rest of the components unchanged.

For voltage clamp recordings, cesium gluconate internal solution was used (composition in mM: 115 cesium hydroxide, 115 gluconic acid, 15 cesium chloride, 2 sodium chloride,10 HEPES, 10 EGTA, 4 Mg-ATP; pH 7.2; 290 mOsm).

#### Data acquisition

Whole-cell recordings were acquired using Multiclamp 700B amplifier, Digidata 1440A digitizer and pCLAMP software (Molecular Devices). The data was low pass filtered at 2 kHz using a Bessel filter and sampled at 50 kHz at a gain of 1. The capacitive currents were compensated using the amplifier circuitry. In case of voltage clamp recordings, leak currents were subtracted. The recordings are not corrected for liquid-liquid junction potential which has earlier been estimated to be +8 mV for the K gluconate based internal solution (Sengupta and Thirumalai, 2015). We measured the neuron’s capacitance, input resistance and series resistance at the start of the experiment. The required capacitance was between 10 and 20 pF and input resistance between 800 MΩ and 3 GΩ. Neurons with resting membrane potential above −30 mV and with series resistance higher than 10% input resistance were excluded. Series resistance was measured at regular intervals during the recordings. Cells in which it had changed by more than 25% in the course of the experiment were excluded.

#### Pharmacology

All pharmacological agents except BAPTA were superfused in the bath. Recordings from the same cell prior to antagonist superfusion were considered as control. Recordings were made after 5-6 minutes of perfusion of the pharmacological agent. The complete list of the chemicals used and their concentrations are given in table 1.

**Table 1:** List of pharmacological agents used for this study.

| Chemical name | Make | Catalog number | Concentration used |
| --- | --- | --- | --- |
| Apamin | Hello Bio | HB1051 | 200 nM |
| Cadmium chloride | Sigma Aldrich | 202908 | 10 μM |
| Gabazine (SR-95531) | Sigma Aldrich | S106 | 10 μM |
| Iberiotoxin | Hello Bio | HB1053 | 100 nM |
| K4-BAPTA | Life Technologies | B1024 | 20 mM |
| Mibefradil | Sigma Aldrich | M5441 | 100 μM |
| NBQX | Tocris | 373 | 20 μM |
| Nickel chloride | Sigma Aldrich | 223387 | 30 μM |
| Nifedipine | Sigma Aldrich | N7634 | 200 μM |
| ω-Agatoxin IVA | Hello Bio | HB1212 | 100 nM |
| TEA Bromide | Sigma Aldrich | 241059 | 1 mM |
| Tetrodotoxin | Hello Bio | HB1035 | 1 μM |

### Analysis

All traces were analyzed and plotted using ClampFit 10.7 and custom scripts written in MATLAB (MathWorks Inc).

#### Estimation of resting membrane potential

To estimate resting membrane potential, the traces were first passed through a moving median filter (window size 120 ms) to remove fast membrane potential changes due to spikes. For the tonic state, resting membrane potential was taken as the mode of membrane potential distribution of the filtered trace. In case of the bursting state, the membrane potential had a bimodal distribution and the two modes were considered as the two resting membrane potentials.

#### Spike detection

The unfiltered traces were used to identify the simple spikes and the climbing fibre inputs. Peaks were first identified based on their prominence from baseline (how much they differ from baseline). In this case, the baseline would be the so called ‘resting membrane potential’ calculated previously. An amplitude threshold was used to distinguish the climbing fiber inputs (> 30 mV) from the simple spikes (10-30 mV). The spike widths of all simple spikes were measured and only spikes with widths between 3-6 ms were included. These values were chosen after manual examination of several cells acquired over the course of this study as well as from cells previously recorded. The rest of the spikes were manually checked and, in most cases excluded.

#### Spike kinetics estimation

The decay time is time required to settle back to 10% of peak amplitude. Action potential width was measured at half amplitude. Threshold was determined to be 4 percent of maximal dv/dt value. AHP values were calculated as the difference between threshold and the most hyperpolarized point immediately following the peak of the actional potential. Bursts were defined as any group of three or more spikes with an interval less than 300 ms.

### Statistics

#### Statistical tests

All statistical tests were implemented using a custom Python script using ‘SciPy (Virtanen et al., 2020). Non-parametric two-sided tests were performed for comparisons as these are best suited best for small sample sizes. Wilcoxon signed-rank test was used for paired samples and Mann-Whitney U for independent samples. Method = ‘Exact’ was used to calculate the exact p-value instead of relying on an approximate normal distribution. P <0.05 was used as a threshold for significance.

#### Statistical Power estimation

Effect sizes were calculated using the rank-biserial correlation. Subsequently, Monte Carlo simulations were performed by generating multiple resampled datasets. For each simulated dataset, the appropriate non-parametric test (Mann-Whitney U or Wilcoxon signed-rank) was conducted to estimate the probabilities. The number of simulations was adaptively determined to minimize the standard error of the estimation, ensuring robust and accurate statistical inference. Effect size, statistical power, number of simulations and standard error of estimations are included in the statistics summary.

p-value and estimated power (α) are mentioned along with all plots. For paired samples, mean ± standard deviations are indicated on respective plots in red. For independent samples, Whiskers on the box plots represent 1.5 times the interquartile range (IQR).

## Author contributions

MPJ: Conceptualization, investigation, analysis and writing; SV: Investigation and analysis; VT: Conceptualization, analysis, and writing.

## Data and materials availability

All data related to this publication is available on Zenodo. DOI 10.5281/zenodo.10222814.

## Acknowledgments

The authors would like to thank the funding agencies listed below and Mr. P.T. Jagadeesh for the maintenance of our fish lines.

